# Dual function of HYPONASTIC LEAVES 1 during early skotomorphogenic growth in Arabidopsis

**DOI:** 10.1101/728527

**Authors:** Juan Manuel Sacnun, Roberta Crespo, Javier Palatnik, Rodolfo Rasia, Nahuel González-Schain

## Abstract

MicroRNAs are small RNA molecules with big impact in many eukaryotic biological processes. In plants, their role as regulators of important developmental programs such as leaf size and shape, flower organs or phase transitions, among others, have been evidenced by mutants in specific miRNAs and by mutants in components of their biogenesis. However, we are still far from understanding the scope of this regulatory system so other crucial developmental phases might be influenced by the microRNA pathway.

Skotomorphogenesis is an essential developmental program that takes place after seeds germinate underground in order to display a proper response when seedlings reach the light. In this work, we found that the core components of microRNA pathway, DCL1, HYL1 and SERRATE, promote hypocotyl elongation during skotomorphogenesis. Hook unfolding, another characteristic phenotype displayed by dark-grown seedlings is also regulated by these proteins but, surprisingly, they act in different ways. Thus, HYL1 represses hook unfolding while DCL1 and SE promote it since the hooks of mutants on each component are more or less open than those of wild-type during skotomorphogenesis, respectively. Genetic and physiological analyses on HYL1 mutants provide evidence that repression of hook unfolding is carried out through the HYL1 protein-protein interaction domain. Furthermore, the data indicates that phosphorylated HYL1 is necessary for this function. Molecular and genetic analyses also suggest that HYL1 regulates the activity of the master photomorphogenic regulator HY5 in darkness to ensure a proper early skotomorphogenic growth. In summary, while our data show a role for miRNAs in darkness, it also suggests a microprocessor-independent role of HYL1 as a repressor of hook unfolding assigning a biological function to phosphorylated HYL1. This work uncovers a previously unnoticed link between components of the miRNA biogenesis machinery, the skotomorphogenic growth and hook development in Arabidopsis.

**Author summary:** Seeds germinating underground display a specific developmental program, termed skotomorphogenesis, to ensure survival of the emerging seedlings until they reach the light. They rapidly elongate the hypocotyl and maintain the cotyledons closed, forming a hook with the hypocotyl in order to protect apical meristematic cells from mechanical damage. Such crucial events for the fate of the seedling are tightly regulated and although some transcriptional regulators and phytohormones are known to be implicated in this regulation, we are still far from a complete understanding of these biological processes. Our work provides new information on the diverse roles in skotomorphogenesis of the core components of microRNA biogenesis in Arabidopsis, HYL1, SE, and DCL1. We show that hypocotyl elongation is promoted by all these components, probably through the action of specific miRNAs. Hook development is also controlled by these components although, remarkably, HYL1 exerts its role in an opposite way to DCL1 and SE. Interestingly, we found that a specific HYL1 domain involved protein-protein interaction is required for this function, instead of other regions of the protein with known roles in the biogenesis of miRNAs. We propose that phosphorylated HYL1 help to maintain the hook closed during early skotomorphogenesis by repressing the activity of HY5, the transcriptional master regulator that triggers light responses.

## Introduction

Seeds buried in the soil have evolved a particular developmental program to grow heterotrophically until the seedling emerges and reaches the light, where it can change its developmental program to autotrophic growth. Thus, growth in darkness, or skotomorphogenesis, is characterized by an exaggerated growth of hypocotyl while cotyledons remain closed, protecting the shoot apical meristem by forming a hook between them and the hypocotyl [1]. Seed reserves are limited, hence a proper skotomorphogenic growth must be achieved in order to ensure survival of seedlings until they reach the light. A complex network of transcription factors and phytohormones such as ethylene, auxins and gibberellins tightly regulates skotomorphogenic growth once seeds germinate and during the transition to light [2, 3]. The bHLH PHYTOCHROME INTERACTING FACTORS (PIFs) exert their promoting role in darkness and, at the same time, repress photomorphogenesis [4]. On the other hand, the bZIP transcription factor ELONGATED HYPOCOTYL5 (HY5) acts as a master transcriptional regulator during photomorphogenesis (revised in [5]). In darkness HY5 is targeted for degradation by the E3 ubiquitin-ligase CONSTITUTIVE PHOTOMORPHOGENIC1 (COP1) [6].

MicroRNAs (miRNAs) are a family of small (21-24 nucleotides) RNAs critical for a number of biological processes in eukaryotes. In plants, these small molecules are able to silence specific target genes by base complementarity and cleavage -or translation inhibition-of target mRNAs. MiRNAs have a great impact in plant growth, development and stress-related responses since many of their target genes encode for transcription factors and other key regulator proteins. They differ from other families of small RNAs in their biogenesis, which is performed by a specific protein complex in the nucleus. DICER LIKE 1 (DCL1), SERRATE (SE) and HYPONASTIC LEAVES1 (HYL1) are considered to be the core complex that process primary messengers of miRNAs (pri-miRNA) into miRNAs. While DCL1 carry out endonucleolytic cleavages of miRNA precursors, SE and HYL1 act as adaptors to ensure the efficiency and accuracy of this process [7–9]. Mutant plants in those proteins display an accumulation of pri-miRNAs and low levels of many families of miRNAs as well as many developmental and growth defects [10–14].

HYL1 comprises two double-stranded RNA binding domains (dsRBD1 and dsRBD2) in its N-terminus end and six consecutive 28 residue repeats of unknown function in its C-terminus. These repeats seem to be expendable for its function, since a truncated version of HYL1 with only the dsRBD1-dsRBD2 region is able to complement *hyl1-2* mutant in leaves [15]. While dsRBD1 is involved in binding to miRNA precursors [16–18], dsRBD2 participates in protein-protein interaction with DCL1 [7, 19] and SE [20], and in homodimerization [16, 21]. HYL1 is also regulated post-translationally by dephosphorylation of serine residues by C-TERMINAL DOMAIN PHOSPHATASE-LIKE 1 (CPL1, [22]), Protein Phosphatase 4 and SMEK complex [23], and is a potential substrate of phosphorylation by MPK3 and SnRK2 kinases [24, 25]. Also, its levels are regulated by degradation by an unknown protease after a transition from light to dark conditions in adult plants [26]. The developmental defects displayed in *HYL1* mutant plants have evidenced the many biological processes in which the microprocessor is involved, such as leaf curvature, elongation of stamen filaments that affect fertility, or altered responses to phytohormones, among others [14].

Some recent studies suggest a role for HYL1 and other components of the microprocessor in early light-grown seedling development and in dark-to-light transition [26–28]. However, the relevance of miRNA biogenesis components during skotomorphogenic growth is still poorly understood. Here we show that the microRNA pathway is required during Arabidopsis skotomorphogenic growth to ensure a proper development of hypocotyls and apical hooks until seedlings reach the light. Our results indicate different roles of core proteins belonging to the microprocessor complex in different tissues during skotomorphogenesis. We found that DCL1, HYL1 and SE promote hypocotyl elongation. As expected, mutants in these genes have similar phenotypes as they act together in miRNA biogenesis. However, we found a more complex scenario in hook development. While DCL1 and SE promote hook unfolding, HYL1 repressed it. The specific role of HYL1 in the repression of hook development seems to operate through protein-protein interactions rather than through miRNA biogenesis. The data point to phosphorylated HYL1, which is inactive for miRNA biogenesis, having a function during normal hook development. We propose a novel regulatory layer linking HYL1 to HY5 during skotomorphogenic growth.

## Results

### HYL1 is necessary for a proper skotomorphogenic growth

During the initial characterization of *HYL1* it was shown that mutant seedlings develop shorter hypocotyls than wild type plants [14, 28]. However, no further analysis was carried out to understand the role of HYL1 on skotomorphogenic development. We tested the main characteristic phenotypes of seedlings grown in darkness, that is hypocotyl elongation and hook development, in time-course experiments from two to four days in darkness (dD). *hyl1-2* mutants have consistently shorter hypocotyls than WT seedlings at all time points analyzed (Figs 1A and 1B). These differences cannot be attributed to a delay in germination of *hyl1-2* seeds since this is not affected (S1 Fig). The requirement of HYL1 to elongate the hypocotyl properly seems to be specific for dark conditions since this mutant did not show any significant differences with WT at early stages in light conditions (in long day, LD; S2A and S2B Figs). The apical hook is significantly more open in *hyl1-2* than in wild-type (WT) seedlings during skotomorphogenic growth (Figs 1A and 1C). Overall, plants lacking HYL1 seem to develop a partial photomorphogenic growth in darkness. Since phenotypic differences can be seen at 2dD and both hypocotyl elongation and hook opening rates remain relatively unchanged at later time points, we hypothesize that the presence of HYL1 is more relevant at early than late stages during skotomorphogenesis.

**Fig 1.**
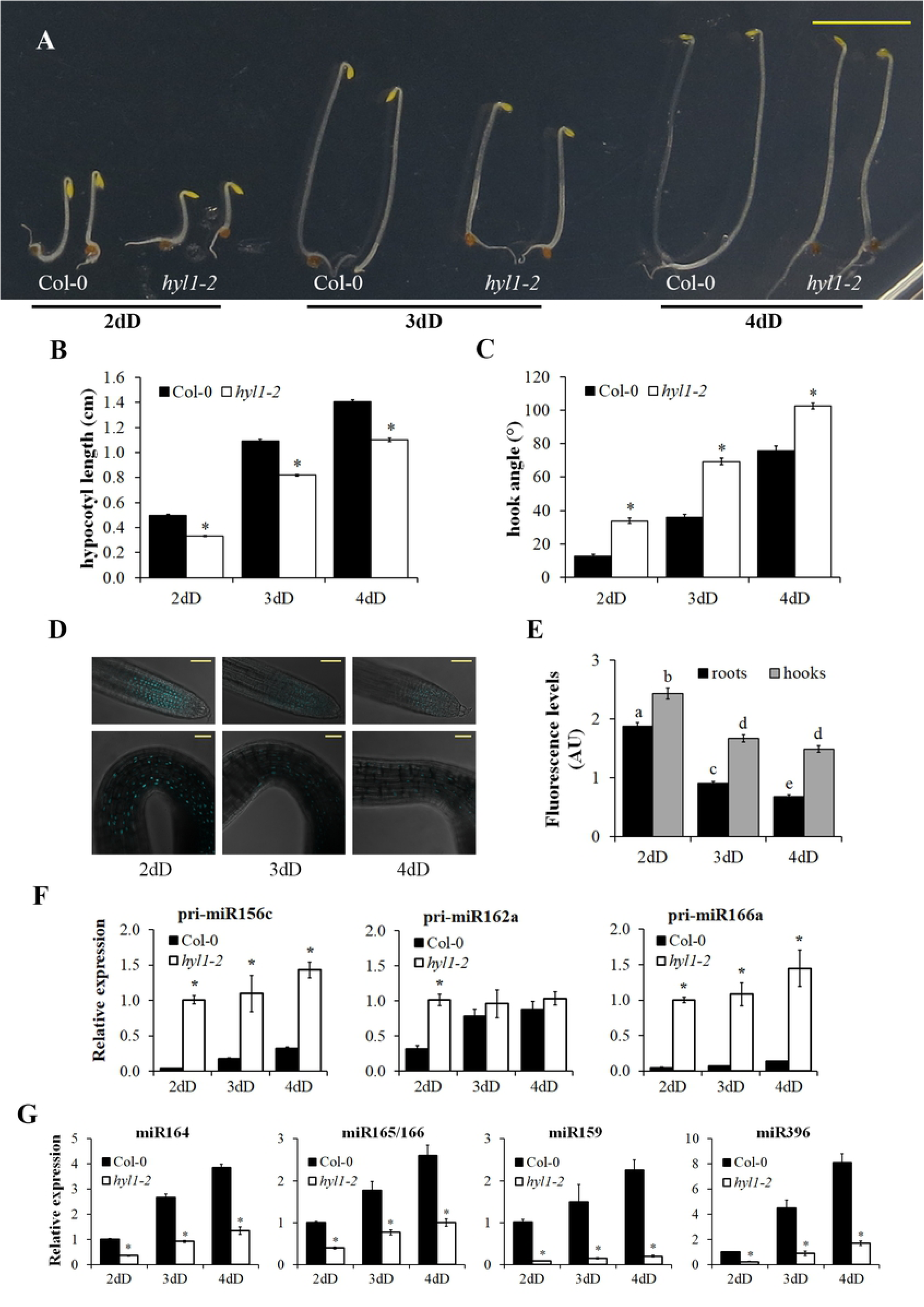
HYL1 is required for early skotomorphogenic growth. (A) Representative *hyl1-2* and control (Col-0) seedlings grown in darkness. dD indicates days in the dark. (B) Hypocotyl length and (C) hook angle measurements of *hyl1-2* and Col-0 dark-grown seedlings at the indicated time points. Data are reported as mean ± SEM of at least 40 seedlings from 5 biological replicas. (D) Representative confocal fluorescence images of roots (upper panels) and hooks (lower panels) from dark-grown pHYL1::HYL1-CFP *hyl1-2* seedlings at the indicated time points. (E) Integrated fluorescence intensity of root and hook nuclei from dark-grown pHYL1::HYL1-CFP *hyl1-2* seedlings at the indicated time points Data are reported as mean ± SEM of at least 70 nuclei from two biological replicas. AU: Arbitrary units. (F) and (G) qRT-PCR and stem-loop qRT-PCR analysis, respectively, of *hyl1-2* and Col-0 dark-grown seedlings at the indicated time points. Expression levels of pri-miRNA (F) or miRNAs (G) were normalized to the *PP2AA3* housekeeping gene and expressed relative to the *hyl1-2* (F) or Col-0 (G) 2dD value set at unity. Means ± SEM are shown from technical triplicates and 2 biological replicas. Asterisks in (B), (C), (F) and (G) indicate statistically different mean values compared with their corresponding wild-type (p < 0.05). Statistically significant differences between groups in (E) are indicated by different letters (ANOVA, p < 0.05). Bar in (A) = 5 mm and in (D) = 50 µm.

To study where is HYL1 actively participating in the control of skotomorphogenic growth we generated *hyl1-2* plants expressing HYL1-CFP under the control of its own promoter (pHYL1::HYL1-CFP; [29]). Several transgenic lines expressing the fusion protein were obtained, and the difference in expression levels between lines was evaluated by measuring fluorescence intensities of nuclei in roots (S3A and S3B Figs). Despite the differences in expression levels, all pHYL1::HYL1-CFP lines were able to complement the phenotypes displayed by *hyl1-2* mutant during skotomorphogenesis (S3C and S3D Figs) and in leaves of adult plants (S3E Fig). HYL1-CFP fluorescence was detected mainly in nuclei of roots, hooks (Fig 1D, S3B and S4A Figs), hypocotyls and cotyledons during skotomorphogenesis. Several seedlings displayed diffuse fluorescence signals in roots (S4B Fig) and other tissues with no clear localization. We chose homozygous lines 10 and 15 with *ca.* 5-fold level differences in HYL1 expression, as quantified by fluorescence, (S3A Fig) for further studies in order to address quantitative aspects of HYL1 localization by confocal fluorescence microscopy. First, we quantified the number of seedlings with discrete nuclear or diffuse localization in different tissues of both lines. A significant proportion of seedlings turn from nuclear to diffuse localization as the skotomorphogenesis progresses in roots and hypocotyls. Interestingly, HYL1-CFP remains nuclear in hooks during all three time points tested in both lines (S1 Table). We then quantified fluorescence intensities of nuclei from those seedlings with discrete nuclear localization. Figs 1D and 1E show that the level of HYL1-CFP decreases in roots as skotomorphogenesis progresses while fluorescence intensities remain constant in hooks after an initial drop from 2dD to 3dD. In summary, our results suggest that HYL1 is present during early stages of development in darkness and is degraded in most tissues as skotomorphogenesis progresses. The particular persistence of HYL1 levels in hooks could indicate that proteolytic regulation is not taking place in that tissue and that the protein could have a role in the repression of hook opening.

HYL1 is a key adaptor within the dicing complex to precisely and efficiently generate microRNAs (miRNAs) from their primary RNA precursors (pri-miRNAs; [7]). Whether this machinery is ubiquitously active or time– and developmentally regulated is not known, and there is currently no information on whether miRNA biogenesis is active during skotomorphogenesis. With a few exceptions [30], plants with mutations in miRNA biogenesis proteins fail to generate miRNAs and accumulate pri-miRNAs. We tested the expression levels of nine pri-miRNAs, 156b, 156c, 156d, 159b, 162a, 164c, 166a, 166b and 171b in *hyl1-2* and Col-0 dark-grown seedlings. All nine precursors accumulate in the mutant at higher levels than in controls at 2dD (Fig 1F and S5 Fig). Moreover, this over-accumulation persists even at 4dD for most of them although with smaller differences. To test whether this accumulation of pri-miRNAs in *hyl1-2* is due to a deficient microprocessor activity we checked the abundance of several miRNAs by stem-loop qPCR. Compared to WT seedlings, a very small amount of miRNAs is produced in *HYL1* mutant seedlings in darkness (Fig 1G). Our findings suggest that HYL1 activity, through the biogenesis of miRNAs, is important during skotomorphogenesis.

### Other components of the microRNA pathway are also implicated in the control of skotomorphogenesis

As HYL1 is actively participating in the control of skotomorphogenesis, and no specific role beside microRNA biogenesis was found for this protein until now, we expected to find a similar role for other components of the microRNA pathway. Together with HYL1, DCL1 and SE form the core component of the microRNA biogenesis machinery. We therefore tested whether *se-1* and *dcl1-100* mutants (both in Col-0 background as *hyl1-2*) displayed similar phenotypes as *hyl1-2* in dark-grown seedlings. Mutants in *SE* exhibited significantly shorter hypocotyls than WT during skotomorphogenesis (Figs 2A and 2B). Unexpectedly, hook angle is less open in *se-1* at all time points tested (Figs 2A and 2C), showing a phenotype opposite to *hyl1-2*. We also tested the behavior of light-grown *se-1* mutants. Although significant differences were found in hypocotyl elongation between *se-1* and WT at 3 days in LD these are rather small and probably with low biological impact (S6A and S6B Figs). This suggests that, as in *HYL1* mutants, the role of SE in early development is more relevant during skotomorphogenesis. In order to study the localization of SE protein in dark-grown seedlings we generated pSE::YFP-SE lines in *se-1* background and checked fluorescence by confocal microscopy. Images obtained from several independent lines grown at 3dD showed localization of YFP-SE in nucleoplasm from all tested tissues: hooks, hypocotyls, roots and cotyledons (Fig 2D). Given that SE is present during skotomorphogenesis and the phenotypes observed in *se-1* we next tested whether miRNA biogenesis is impaired in dark-grown mutant seedlings. We tested the expression of nine pri-miRNAs in *se-*1 and WT seedlings grown 2dD and 4dD by qPCR. Results showed an overaccumulation of all pri-miRNAs tested in *se-1* (Fig 2E and S5 Fig) indicating miRNA biogenesis is active during skotomorphogenesis, as was suggested previously from *hyl1-2* experiments. Similar to what we observed in *hyl1-2* mutants, miRNAs accumulation is also impaired in *se-1* during skotomorphogenesis (Fig 2F). We tested for phenotypic alterations in dark-grown *DCL1* mutants as well, by using the hypomorphic allele *dcl1-100*. As homozygous mutants are sterile we carried out experiments in darkness using *dcl1-100* heterozygous lines and its WT siblings as controls. Dark-grown homozygous *dcl1-100* seedlings are easily recognized in a segregating population in the plates after four more days growing in light (S7A and S7B Figs). Their cotyledons are hardly completely opened, as in *hyl1-2* mutants (S2A Fig), and segregation analyses correlate well with genotypes of these seedlings (S2 Table). Hypocotyl elongation rate is significantly slower in *dcl1-100* compared to the rest of heterozygous and WT segregating seedlings, as well as to WT siblings grown in parallel (S7A and S7C Figs), thus resembling *hyl1-2* phenotypes. In contrast, hook angle is less open in *dcl1-100* at all time points tested (S7A and S7D Figs), this phenotype being the same as the *SE* mutants and opposite to *hyl1-2*. Taken together, our results demonstrate that miRNA biogenesis is required to rapidly elongate the hypocotyl seeking light during skotomorphogenesis. Hook development is also controlled by core components of miRNA pathway, but our results identify a specific role for HYL1 that acts in an opposite way to DCL1 and SE. This led us to study the role of HYL1 on skotomorphogenesis in more detail.

**Fig 2.**
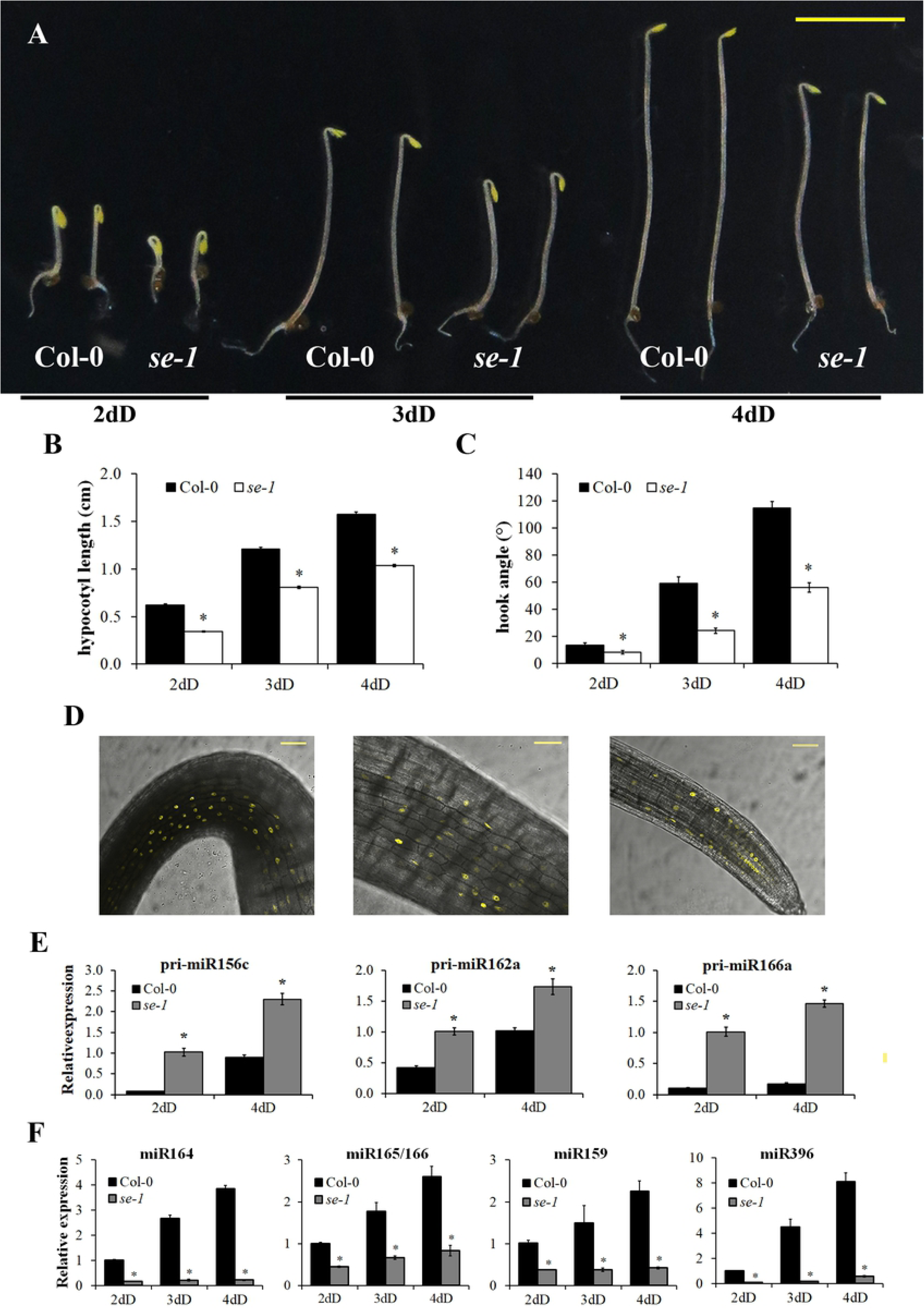
SE participation in skotomorphogenesis. (A) Representative *se-1* and control (Col-0) seedlings grown in darkness are shown. dD indicates days in the dark. (B) Hypocotyl length and (C) hook angle measurements of *se-1* and Col-0 dark-grown seedlings at the indicated time points. Data are presented as mean ± SEM of at least 40 seedlings from 2 biological replicas. (D) Representative confocal fluorescence images of hooks, hypocotyls and roots from 3-days dark-grown T1 pSE::YFP-SE seedlings. (E) and (F) qRT-PCR and stem-loop qRT-PCR analysis, respectively, of *se-1* and Col-0 dark-grown seedlings at the indicated time points. Expression levels of pri-miRNAs (E) or miRNAs (F) were normalized to the *PP2AA3* housekeeping gene and expressed relative to the *se-1* (E) or Col-0 (F) 2dD value set at unity. Means ± SEM are shown from technical triplicates and 2 biological replicas. Asterisks in (B), (C), (E) and (F) indicate statistically different mean values compared with their corresponding wild-type (p < 0.05). Bar in (A) = 5 mm and in (D) = 50 µm.

### HYL1 dsRBD1 is required for miRNA biogenesis but not for hook development during skotomorphogenesis

HYL1 comprises two double-stranded RNA-binding domains (dsRBDs) in its N-terminal end, a nuclear localization signal and six repetitions of 28 aminoacids in its C-terminal region (Fig 3A). In order to dissect the role of particular domains of HYL1 during skotomorphogenesis we tested three mutant transgenic lines in dsRBD1. We have previously demonstrated that mutations in dsRNA binding region 1 are most detrimental to the activity of HYL1 *in vivo*, whereas mutations in regions 2 and 3 do affect its activity to a lesser extent [18]. Overexpression of HYL1 mutants in region 1 of dsRBD1 (mR1: K17A/R19A) did not complement *hyl1-2* hypocotyl elongation, while a seven-aminoacids deletion Δ40-46 in region 2 (mR2) and region 3 mutants (mR3: R67A/K68A) only partially complemented the hypocotyl elongation (Figs 3B and 3C, and S8 Fig). The requirement of dsRBD1 HYL1 in hypocotyl elongation goes in line with its role in miRNA biogenesis and function in leaf development [18]. Surprisingly, all mR1, mR2 and mR3 mutants almost fully complemented hook phenotypes in *hyl1-2* (Figs 3B and 3D). These results suggest that dsRBD1 of HYL1 is involved in the control of hypocotyl elongation as well as in leaf curvature, but does not participate in the regulation of hook development during skotomorphogenesis. Furthermore, it hints that the participation of HYL1 in hook development is independent of its role in miRNA biogenesis, as pri-miRNA binding by HYL1 is dominated by dsRBD1 [17–19].

**Fig 3.**
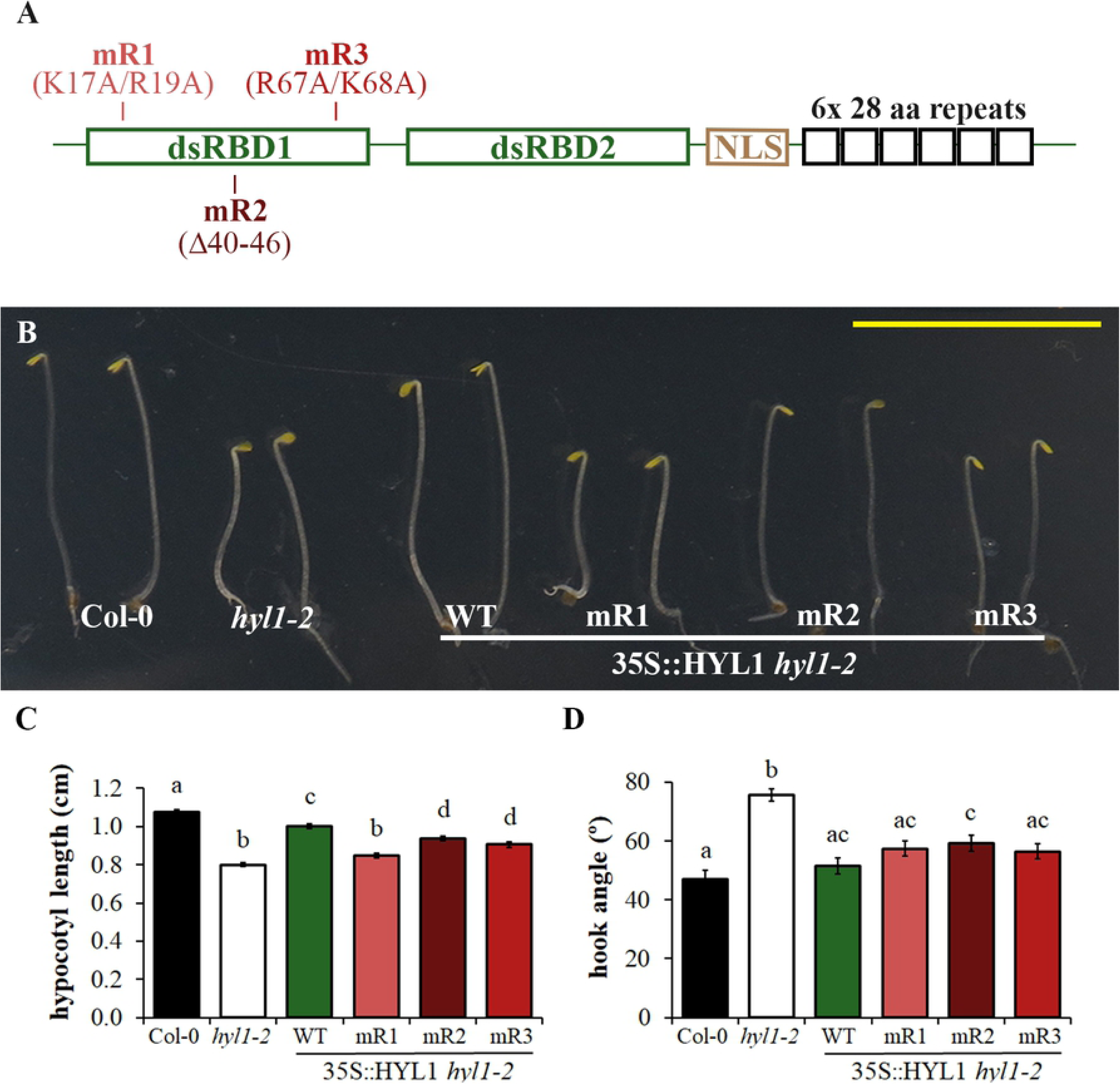
The dsRBD1 domain of HYL1 is not involved in hook unfolding regulation in darkness. (A) Schematic view of HYL1 protein. The two double-stranded RNA-binding domains (dsRBD), nuclear localization signal (NLS) and repetitions at the C-terminal end are indicated. mR1, mR2 and mR3 mutant versions in dsRBD1 [18] are also indicated. (B) Representative *hyl1-2*, control (Col-0) and plants expressing wild-type or mutant versions of HYL1 dsRBD1 in *hyl1-2* background from 3-days dark-grown seedlings are shown. dD indicates days in the dark. (C) Hypocotyl length and (D) hook angle measurements of the plants shown in (B). Data are presented as mean ± SEM of at least 40 seedlings from 2 biological replicas. Statistically significant differences between groups in (C) and (D) are indicated by different letters (ANOVA, p < 0.05). Bar in (B) = 5 mm.

### Hook unfolding is repressed in mutant lines that affect the phosphorylation status of HYL1

The activity of HYL1 is regulated by phosphorylation in specific serine residues. It was shown that phosphorylated HYL1 displays a reduced activity and that recruitment to the microprocessor complex is impaired leading to defects in miRNA processing and accumulation [22]. In order to verify if phosphorylation regulates HYL1 activity in skotomorphogenesis as well, we analyzed the behavior of mutants that modify the phosphorylation state of HYL1. C-terminal domain phosphatase-like 1 (CPL1) is the main phosphatase that reactivates HYL1 by dephosphorylation of serine residues [22]. We tested a strong (*cpl1-7*) and a mild (*cpl1-3*) allele of *CPL1* mutants during skotomorphogenesis. Both *cpl1* mutants displayed only small differences in hypocotyl elongation (Fig 4A). In contrast, hook unfolding is more repressed in both *cpl1-3* and *cpl1-7* than in WT plants (Fig 4B). The strength of *cpl1-3* and *cpl1-7* phenotypes in hooks correlates well with size defects observed in adult plants (S9 Fig). As *CPL* mutants overaccumulate hyperphosphorylated HYL1 [22], their hook phenotypes suggest an active role for phosphorylated HYL1 in the repression of hook unfolding during darkness. In order to confirm that hook phenotypes in *CPL1* are mediated by HYL1 we made double mutants *hyl1-2 cpl1-3* and *hyl1-2 cpl1-7* and tested them in darkness. Hook unfolding measurements showed *hyl1-2* was epistatic to *cpl1-3* or *cpl1-7* as double mutants phenocopied *hyl1-2* in 2-days dark-grown seedlings (S10 Fig). However, this genetic interaction fades away progressively as *hyl1-2 cpl1-3* or *hyl1-2 cpl1-7* resemble wild-type hooks in 4dD (S10 Fig), in coincidence with the early time-window activity of HYL1 in hooks proposed previously.

**Fig 4.**
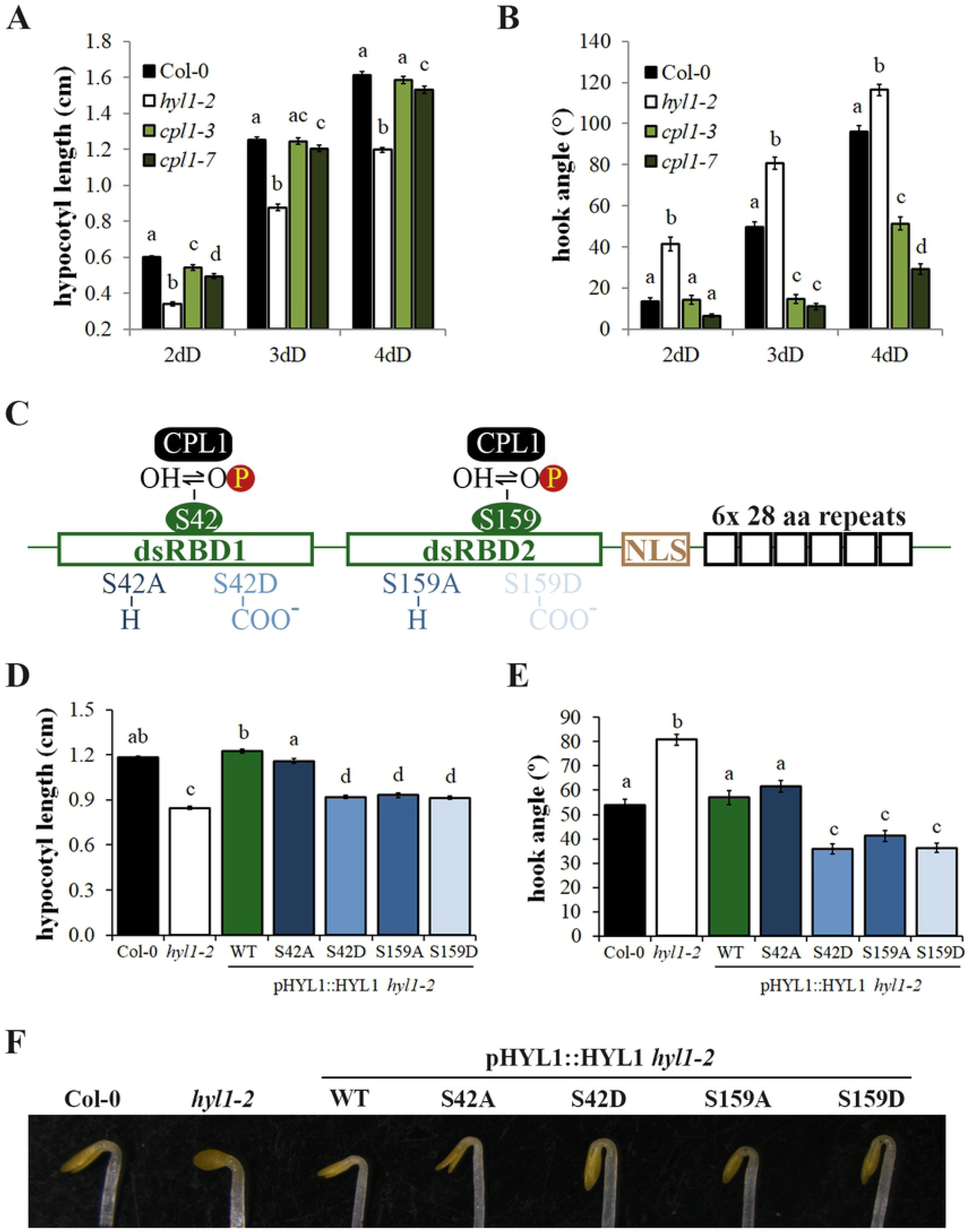
Mutant lines that affect phosphorylation of HYL1 display more closed hooks. (A) Hypocotyl length and (B) hook angle measurements of *hyl1-2*, Col-0 and two mutant alleles of *CPL1* from 2, 3 and 4-days dark-grown seedlings. (C) Schematic view of HYL1. Serines 42 and 159 subjected to dephosphorylation by CPL1 are indicated. Hypo and hyperphosphorylation mimics (S42A, S159A and S42D, S159D, respectively; [22]) are also indicated. (D) Hypocotyl length and (E) hook angle measurements of *hyl1-2*, Col-0, and plants expressing wild-type version or hypo/hyperphosphorylation mimics in *hyl1-2* background from 3-days dark-grown seedlings. (F) Representative hooks of transgenic lines from (E). Data are presented as mean ± SEM of at least 40 seedlings from 2 biological replicas. Statistically significant differences between groups in (A), (B), (D) and (E) are indicated by different letters (ANOVA, p < 0.05).

Two specific serine residues were described as important for HYL1 phosphorylation mediated regulation: S42 in dsRBD1 and S159 in dsRBD2 (Fig 4C). Mutant versions of S42 or S159 to aspartic acid (S42D or S159D), mimicking constitutive phosphorylation, failed to complement *hyl1-2* defects in leaves, while mutants of seven serines, including S42 and S159, to alanine produced hypophosphorylation mimics of HYL1 that fully restore *hyl1-2* defects [22]. We tested whether these mutant lines were able to complement the defects observed in *hyl1-2* during skotomorphogenesis. Hyperphosphorylation mimic versions of HYL1 S42D and S159D were not able to complement the shorter hypocotyls of *hyl1-2* (Fig 4D) nor the leaves phenotypes of adult plants (S11 Fig). In contrast, S42D and S159D mimics displayed more closed hooks compared to the WT version or Col-0 at 3dD (Fig 4E and 4F). These results further confirm the activity of phosphorylated HYL1 in the repression of hook unfolding.

We also tested the hypophosphorylated mutant versions of HYL1, S42A and S159A. The S42A mutant almost fully complemented both the hypocotyl phenotype and the defects in hook opening of *hyl1-2* (Fig 4D). In line with this, the same mutant restored the curvature in leaves (S11 Fig), confirming that the mutation keeps HYL1 active. In contrast, complementation by the S159A mutant in hypocotyls was virtually insignificant (Fig 4D), and the mutant form also leads to more closed hooks (Fig 4E and 4F). HYL1-S159A shows only partial complementation of the leaf phenotype (S11 Fig), suggesting that the activity of the second dsRBD of HYL1 is partially impaired by the mutation.

### HYL1 controls the activity and stability of HY5 during skotomorphogenesis

Given the skotomorphogenic phenotypes displayed in *hyl1-2* and *se-1* mutants, we suspected that dark/light signaling responses could be altered in these mutants. PIFs transcription factors are essential to maintain skotomorphogenic growth by repressing photomorphogenic responses at a transcriptional level in darkness [4]. *PIL1* and *XTR7* are targets of PIFs and considered as reporters of PIF transcriptional activity [31, 32]. We measured the relative expression of *PIF3*, one of the main components of PIF family with an active role in darkness, and the two targets mentioned above by real-time PCR. Although *PIF3* expression show statistically significant differences between *hyl1-2* and *se-1* mutants, these are rather small and no differences were detected between both mutants and WT (Fig 5A and 5B). These small differences do not seem to reflect a differential transcriptional activity of PIFs between the mutants since no differences in *PIL1* or *XTR7* expression were detected (except for slightly higher levels of *PIL1* expression in *hyl1-2* at 4dD) (Fig 5B). Taken together, neither the changes in PIFs levels nor their activity in *hyl1-2* or *se-1* mutants justify the partial photomorphogenic growth observed in darkness.

**Fig 5.**
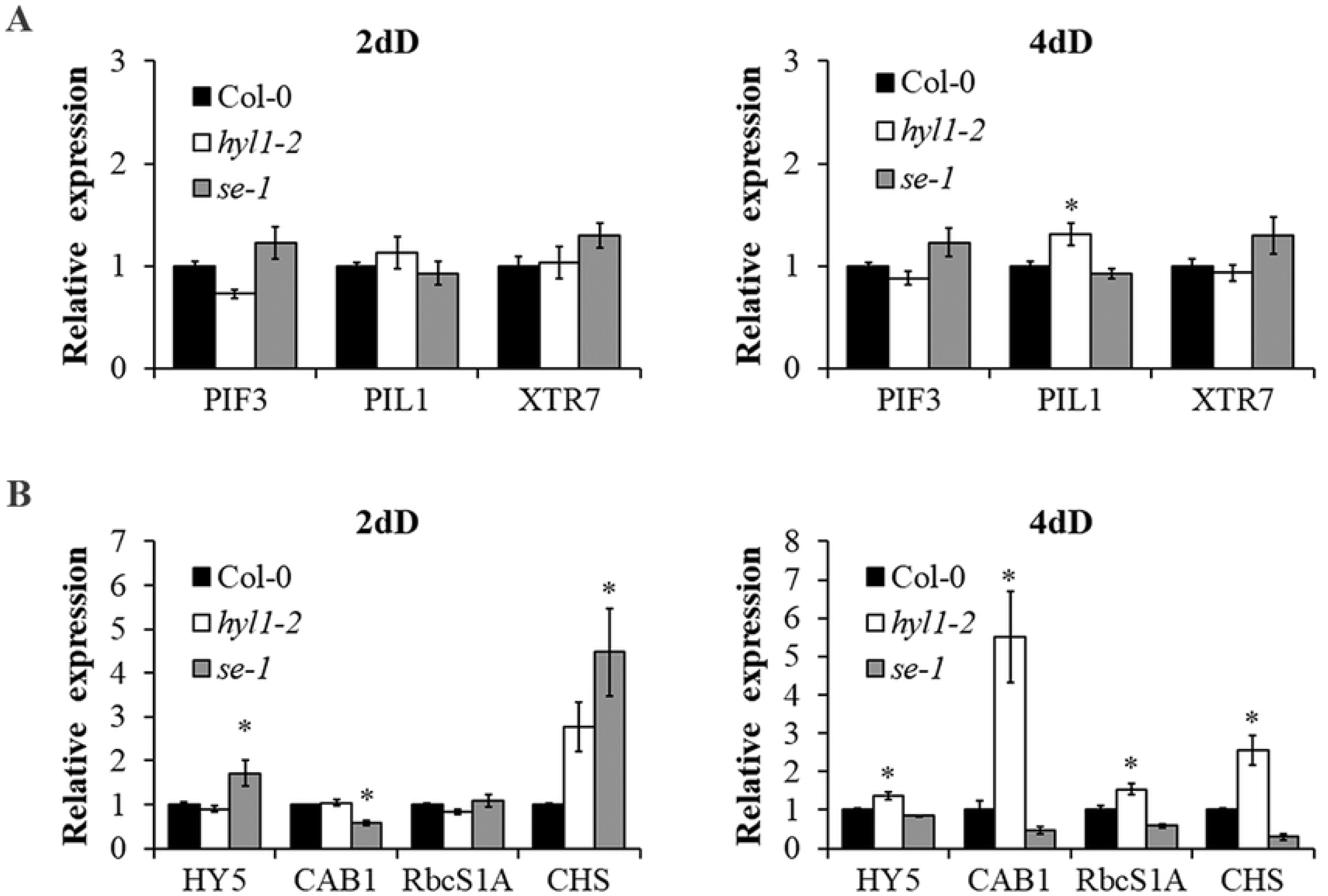
Photomorphogenesis marker genes are repressed by HYL1 in darkness. Expression analysis of (A) *PIF3* and; (B) *HY5*, and their corresponding direct target genes at 2dD (left) and 4dD (right). qRT-PCR analysis of *hyl1-2*, *se-1* and Col-0 dark-grown seedlings at the indicated time points were performed. Expression levels were normalized to the *PP2AA3* housekeeping gene and expressed relative to the Col-0 value set at unity. Means ± SEM are shown from technical triplicates and 2 biological replicas. Asterisks indicate statistically different mean values compared with their corresponding wild-type (p < 0.05).

HY5 transcription factor plays a crucial role in photomorphogenic growth promoting expression of light-regulated genes by binding directly to their promoters [33, 34]. It was shown that HY5 is necessary for the transcription of *Chlorophyll A/b Binding protein1* (*CAB1*), *Ribulose bisphosphate carboxylase Small subunit 1A* (*RbcS1A*) and *CHalcone Synthase* (*CHS*) during the dark-to-light transition [34]. Therefore, we examined the expression levels of these three genes and *HY5* in dark-grown *HYL1* and *SE* mutants by real-time PCR. Statistically significant changes in expression of *HY5*, *CAB1* and *CHS* in *se-1* compared to Col-0 were detected at 2dD (Fig 5C). Though small, these differences cannot be easily explained because a small induction of *HY5* in *se-1* at 2dD was not accompanied with an induction of its direct targets tested. On the other hand, results consistently show a higher expression level of *HY5* in *hyl1-2* that is in line with the induction of *CAB1*, *RbcS1A* and *CHS* observed at 4dD (Fig 5D). Taken together, these results suggest that a small fraction of HY5 protein might be active in *hyl1-2* mutants in darkness partially explaining physiological phenotypes observed during skotomorphogenesis.

In order to test this hypothesis, we made double mutants of *HY5* and *HYL1* or *SE* and measured their hypocotyl elongation and hook unfolding during skotomorphogenesis. *hy5-2* mutants did not show any phenotypic differences compared to WT seedlings, neither in hypocotyl length nor in hook angle, most probably because Constitutive Photomorphogenic 1 (COP1) targets HY5 for proteasome-mediated degradation in the dark [6]. Hypocotyl elongation measurements showed *hyl1-2* or *se-1* were epistatic to *hy5-2*, as both *hyl1-2 hy5-2* and *se-1 hy5-2* phenocopied *hyl1-2* and *se-1* defects, respectively (Fig 6A). A similar epistatic interaction is shown for *se-1 hy5-2* in hook unfolding where the closed hook phenotype in *se-1* was not affected by the absence of HY5 (Fig 6B and 6C). However, we observed different effects when we look at the hook phenotypes. Interestingly, we found that the *hy5-2* mutation is able to restore hook phenotypes when introduced in the *hyl1-2* mutant seedlings (Fig 6B and 6C), suggesting again that *HY5* misregulation in *hyl1-2* mutant was responsible for the hook defects.

**Fig 6.**
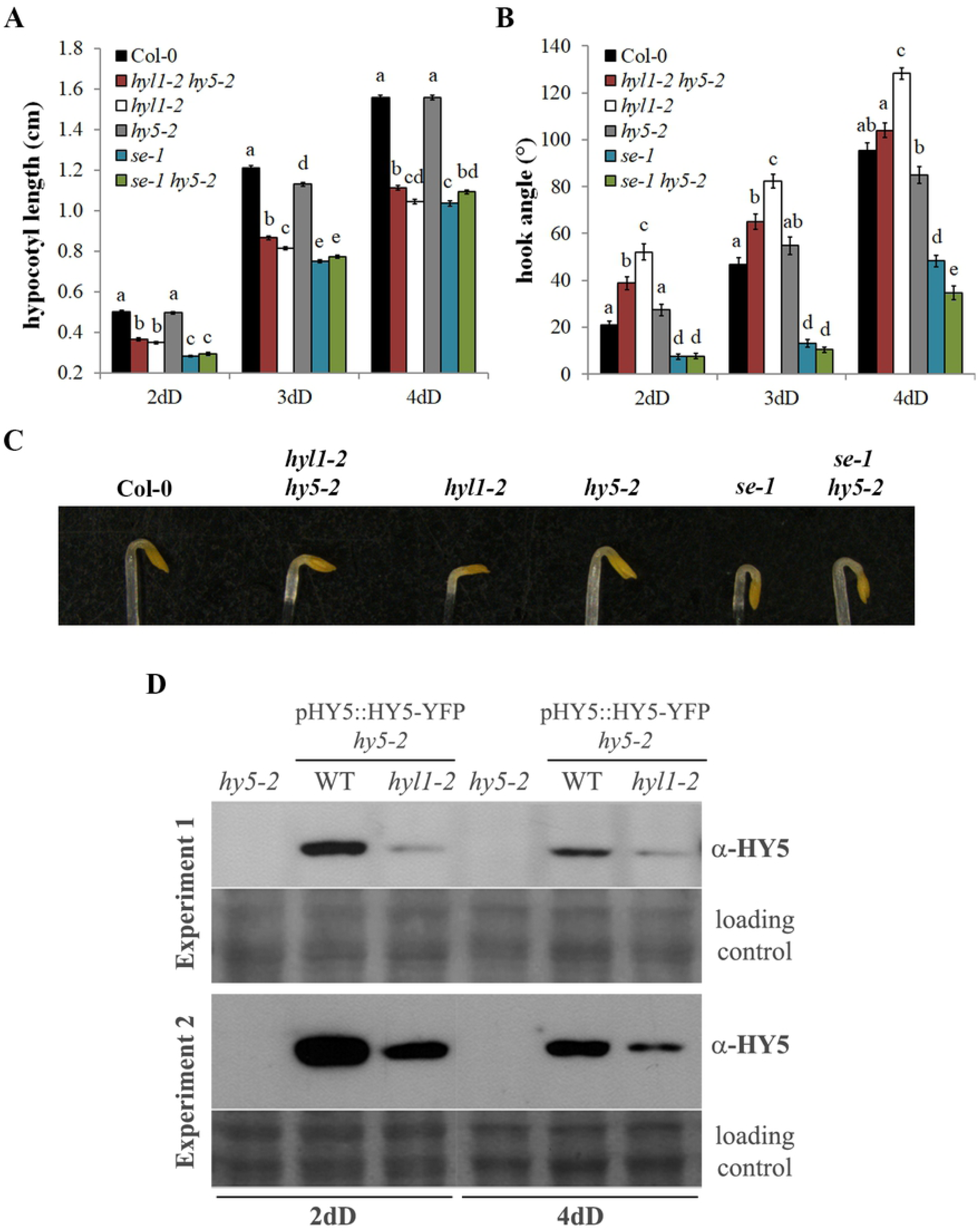
Suppression of *HY5* partially restores hook defects in *HYL1* mutants. (A) Hypocotyl length and (B) hook angle measurements of Col-0, *hyl1-2*, *se-1*, *hy5-2*, and double mutants *hyl1-2 hy5-2* and *se-1 hy5-2* dark-grown seedlings at the indicated time points. (C) Representative hooks of transgenic lines from (B). (D) α-HY5 western blots of 2 and 4-days dark-grown pHY5::HY5-YFP *hy5-2* in *hyl1-2* or Col-0 background seedlings. *hy5-2* mutants were used as negative controls. Ponceau stainings were used as loading controls. Data in (A) and (B) are presented as means ± SEM of at least 40 seedlings from 2 biological replicas. Statistically significant differences between groups are indicated by different letters (ANOVA, p < 0.05).

As HY5 levels in darkness were reported to be very low and inactive due to phosphorylation and proteasome-mediated degradation ([6, 35]), we measured HY5 levels in pHY5::HY5-YFP *hy5-2* and pHY5::HY5-YFP *hy5-2 hyl1-2* by western blot. HY5 is less abundant in *hyl1-2* background at both time-points analyzed (Fig 6D). To exclude a possible transcriptional effect, we analyzed the expression of *HY5* in both lines by qPCR and we found no significant differences (S12 Fig). These results show that HYL1 controls HY5 stability in skotomorphogenesis.

## Discussion

### miRNA biogenesis pathway promotes hypocotyl elongation in skotomorphogenesis

Earlier reports have shown that *Arabidopsis thaliana* mutants in *HYL1* and *DCL1* develop shorter hypocotyls than WT plants at specific time points in darkness [14, 27, 28] but the consistency of this phenotype during skotomorphogenesis has not been previously addressed. Our results indicate that proteins belonging to the core miRNA processing complex promote hypocotyl elongation in darkness since individual mutants in all three major components of this pathway display shorter hypocotyls. This regulation seems to be specific for skotomorphogenic growth since hypocotyl elongation in diurnal conditions is not affected in *hyl1-2* or *se-1* mutants, as was shown in a very recent report for *HYL1* mutants [27]. The complementation of hypocotyl elongation of *hyl1-2* mutants with versions of HYL1 bearing alterations in the pri-miRNA interaction domain (dsRBD1) parallel the complementation of phenotypes in adult plants. These results reveal that specific miRNAs might control cell elongation in hypocotyls. Expression analyses by real-time PCR clearly showed that miRNA biogenesis is actually taking place during skotomorphogenesis. Both miRNA and pri-miRNA levels increase during development in darkness. The more likely explanation for the increase in levels of miRNA precursors is that microprocessing fades out during skotomorphogenesis and that miRNA biogenesis is more relevant at early stages. This agrees with the decay of HYL1 levels through skotomorphogenic development evidenced by quantitative data obtained by fluorescence microscopy. The decreasing relevance of HYL1 during skotomorphogenesis is also evidenced on the dominant phenotypes of *hyl1-2* in double mutants with *cpl1-3* or *cpl1-7*. Hook unfolding is derepressed at early stages but not at later stages, similar to effects observed in *hyl1-2 hy5-2* double mutants. A recent report showed that some miRNAs, like miR319b, miR160b, miR167b and miR848, act as positive or negative regulators of hypocotyl elongation in seedlings grown under red light, although they are not involved in the same process in darkness [27]. Further studies on candidate miRNA mutants or overexpressors would shed light on which are the main miRNA that control hypocotyl elongation during skotomorphogenesis.

### Hook unfolding in darkness is transiently repressed by HYL1

A mutant allele of *HYL1* in Nossen ecotype (*hyl1-1*) was shown to display more open hooks than WT at 3 days in darkness [14], but this finding was not further explored. The data presented here indicates that hook unfolding is also controlled by the components of the microRNA pathway but, intriguingly, while HYL1 is necessary to repress it, SE and DCL1 perform an opposite role promoting hook aperture. Our results also showed that the first dsRBD of HYL1 has a minor role in the control of hook unfolding since complementation of *hyl1-2* phenotypes in hook opening and adult leaves, by *HYL1* mutants in this domain, are clearly dissociated. Although complementation of *hyl1-2* with HYL1-S42D mutants gives a hook phenotype, this serine seems to be dispensable to complement *hyl1-2* hooks because the mR2 mutant, in which a loop containing S42 has been deleted, behaves as WT. On the other hand, *hyl1-2* complementation with phosphomimics points to dsRBD2, the protein-protein interaction domain [7, 16, 19–21], as the defining region in the control of hook unfolding. Taken together, the present work constitutes the first report showing a role for HYL1 that is not linked to miRNA processing.

Our main hypothesis is that phosphorylated HYL1 represses hook unfolding in early skotomorphogenesis, peaking soon after seeds germinate to ensure protection of the meristem until seedlings reach the light. A number of findings support this claim: 1-Complementation of *hyl1-2* mutants with hyperphosphorylation mimics S42D and S159D results in hooks more closed than WT. 2-The same phenotypes were displayed by *se-1* and two alleles of *cpl1* mutants, where only the phosphorylated pool of HYL1 can be detected [22, 36]. 3-A recent report showed an almost exclusive population of phosphorylated HYL1 in darkness, including 4-days old dark-grown seedling [36]. 4-Hyperphosphorylated HYL1 was detected in nucleoplasm but it was not found in dicing bodies [22]. We want to highlight the specificity of phosphorylated HYL1 action during skotomorphogenesis as it was previously claimed that this population is inactive during extended periods of light deprivation [36]. It is worth mentioning that the independent action of HYL1 in hypocotyl elongation and hook unfolding might be related to its stability in the different tissues, as shown in Fig 1E and S1 Table. This can also be the reason for the relevance of CPL1 in hook unfolding, that contrasts with the minor role it plays in hypocotyl elongation. Recently, post-translational modification of HYL1 by CPL1 was shown to be affected by REGULATOR OF CBF GENE EXPRESSION 3 (RCF3) specifically in apex-enriched tissues [37]. We can envisage a similar scenario for hooks and hypocotyls during skotomorphogenesis although more work in specific cell-types is needed to produce a complete picture of this regulation.

### HYL1 controls skotomorphogenesis partly through regulation of HY5 activity

HY5 is a master transcriptional regulator in the de-etiolation process of dark-grown seedlings to promote photomorphogenesis (revised in [5]). As resources from the seeds should be well administrated until seedlings reach the light, HY5 activity must be completely shut-down in darkness to avoid triggering of useless biological processes. It was shown that unphosphorylated HY5, the active form of this transcription factor, is ubiquitinated by COP1 in the nucleus and subject to proteasome-mediated degradation in darkness [6, 35]. Nevertheless, a small pool of phosphorylated HY5 protein, a form less physiologically active and more resistant to degradation, is still present in darkness [35]. This population was postulated to be necessary to quickly respond to illumination during the transition from dark to light [35]. Thus, activity and stability of HY5 are regulated in an opposite direction in darkness and largely depend on its phosphorylation status. Our data clearly shows that HY5 stability is compromised in *HYL1* mutants during skotomorphogenesis. Although much effort is needed to understand the mechanism of this interaction, one plausible explanation is that the phosphorylation status of HY5 might be displaced to a more active population in *HYL1* mutants. Molecular and genetic data presented in this work support this hypothesis. First, we show that at least three direct targets of HY5 are overexpressed in dark-grown *hyl1-2* mutants. Second, epistasis analyses in double and single mutants of *HY5* and *HYL1* strongly suggest HYL1 negatively regulates HY5 activity in darkness. *hyl1-2 hy5-2* double mutants showed a hook opening dynamics intermediate to single mutants or, in other terms, the absence of HY5 partially restored *hyl1-2* hook phenotype. The presence of an active population of HY5 acting in *hyl1-2* mutants could explain its partial photomorphogenic growth in darkness. Full complementation was not achieved probably due to the presence of HY5 homolog (HYH) in *hyl1-2 hy5-2*. HYH showed functional redundancy in other developmental processes such as hypocotyl growth, lateral root growth or pigment accumulation in light (revised in [5]). Also, dynamics of complementation can be explained, as was suggested before, by a repressive role of phosphorylated HYL1 after seeds germinate that fades out during skotomorphogenesis. Further research will reveal if the role of HYL1 on HY5 activity is direct or if it is indirectly exerted by COP1. Although *HY5* mRNA was shown to be target of miR157d in light conditions ([28]), our work suggest that this regulation is not taking place in darkness since expression analyses showed little or no differences in *HY5* expression levels in *hyl1-2* or *se-1* compared to WT.

In summary, this work provides evidence that the microprocessor adaptor HYL1 controls important steps in seedling establishment during skotomorphogenesis in order to successfully develop from heterotrophic to autotrophic growth. Our results strongly suggest that HYL1 in its phosphorylated form have a role in the control of hook development in darkness. We demonstrate that miRNA biogenesis is required for hypocotyl elongation in order to reach the light during skotomorphogenic growth as well. Finally, tight control of the activity of HY5 in darkness could add a new layer of regulation mediated by HYL1, although further research is needed to elucidate whether this process is COP1 dependent or independent. Several questions emerge from this study, which will be uncovered in the future by addressing the mechanism of this new role for HYL1 and how other HYL1-interacting factors influence its response.

## Material and Methods

### Plant material and growth conditions

All mutants used in this work are in the Col-0 background. Mutant plants *hyl1-2* (SALK_064863) and *se-1* (CS3257) were obtained from ABRC (Arabidopsis Biological Resource Center) and have been described elsewhere [11, 13]. Transgenic lines p35S::HYL1 WT, p35S::HYL1 K17A/R19A (mR1), p35S::HYL1 Δ40-46 (mR2) and p35S::HYL1 R67A/K68A (mR3) in *hyl1-2* background were described elsewhere [18]. Transgenic lines pHYL1::HYL1 WT, pHYL1::HYL1 S42A, pHYL1::HYL1 S42D, pHYL1::HYL1 S159A, pHYL1::HYL1 S159D in *hyl1-2* background, *cpl1-3*, *cpl1-7* [22], and *dcl1-100* mutant lines [38] were kindly provided by Dr. Manavella. *hy5-2* (SALK_056405) mutant lines was a kind gift from Dr. Martínez-García. pHY5::HY5-YFP *hy5* was kindly provided by Dr. Botto. *hyl1-2 hy5-2* and *se-1 hy5-2* double mutants were obtained by crossing single homozygous mutants to obtain F2 progenies that were genotyped by PCR-amplification of genomic DNA with specific primers (S3 Table) as follows: hy5-2 gntyp F and hy5-2 gntyp R primers were used to amplify a band of ∼250 bp corresponding to wild-type chromosome; LbB1 and hy5-2 gntyp were used to amplify a band of ∼240 bp corresponding to the chromosome containing the *HY5* disruption by the T-DNA; se-1 gntyp F and se-1 gntyp R were used to amplify a band of 114 bp or 107 bp, corresponding to wild-type or the 7 bp deletion of *SE* containing chromosomes, respectively. Purified bands from agarose gels were further digested with the restriction enzyme SpeI, which results in two fragments of 77 bp and 30 bp only for *se-1* mutation. *hyl1-2* mutation was followed by selection on plates with 50 mg/l Kanamycin due to the T-DNA insertion near the translation start site. *hyl1-2 cpl1-3* and *hyl1-2 cpl1-7* double mutants were obtained by crossing single homozygous mutants to obtain F2 progenies that were genotyped by PCR-amplification of genomic DNA with specific primers (S3 Table) as follows: cpl1-3 gntyp F and cpl1-3 gntyp R primers were used to amplify a band of 341 bp. Purified bands from agarose gels were further digested with the restriction enzyme HindIII, which results in two fragments of 311 bp and 30 bp only for *cpl1-3* mutation; cpl1-7 gntyp F and cpl1-7 gntyp R primers were used to amplify a band of 102 bp. Purified bands from agarose gels were further digested with the restriction enzyme PstI, which results in two fragments of 76 bp and 26 bp only for WT.

Plants in soil and seedlings in plates were grown in growth chambers in long-day (LD) photoperiod (16 h light/8 h dark, ∼100 μmol m^-2^ sec^-1^) or constant darkness at 22°C.

### Generation of transgenic lines

Plasmids containing pHYL1::HYL1-CFP and pSE::YFP-SE ([29]) used for transformation of *hyl1-2* or *se-1* plants, respectively, were a kind gift from Dr. Fang and Dr. Spector. *Agrobacterium tumefaciens* strain GV3101 containing each construct with the respective translational fusion under the endogenous promoter were obtained and used to transform mutant plants by floral dip method [39]. Selection of each transgenic line in the following generations were performed as described [29].

### Seedling growth and measurements

Seeds were surface-sterilized for 2 min in 96% EtOH, transferred to 20% v/v NaOCl and 0.05% v/v triton X-100 for 10 min, and rinsed five times with sterile distilled water. Seeds were then placed in petri dishes containing half-strength Murashige and Skoog (MS) agar medium, covered with aluminum foil and stratified for 5 days in darkness at 4°C. Germination was induced with 3h of white light in the growth chamber, plates were then covered with aluminum foil and placed inside a box, and incubated in the same growth chamber for the time points indicated in each experiment. For hypocotyl elongation and hook angle measurements, seedlings were arranged horizontally on a plate, photographed using a digital camera (Canon PowerShot SX50 HS) and measured with ImageJ 1.49k software. Hook angle was measured as the angle between the hypocotyl and an imaginary line between the cotyledons.

Experiments with *dcl1-100* seeds were carried out as described above but arrangement of dark-grown seedlings on the plate and photographs were performed in sterile conditions in a hood. Plates were then incubated for 4 more days in LD and photographs were taken. Defective and normal seedlings (S6D Fig) were identified from the photographs of light-grown seedlings and used to mark and do the measurements from the photographs of dark-grown seedlings. Segregation analysis was performed in order to confirm genotypes of mutant seedlings (S2 Table).

### Confocal microscopy

Dark-grown seedlings of pHYL1::HYL1-CFP *hyl1-2* and pSE::YFP-SE *se-1*, at indicated times for each experiment, were transferred to glass slides, mounted in water and covered with a coverslip. Laser confocal microscopy was performed using a Plan Apochromat 20X, 0.8 NA lens on a ZEISS LSM880 microscope. Fluorescent fusion proteins were excited with the 458 nm (for CFP) and the 514 nm (for YFP) lines of an argon laser and fluorescence emission was collected from 459-522nm (CFP) or 517-570 nm (YFP). Pictures were taken and fluorescence signal intensity of individual nucleus were analyzed with ImageJ 1.49k software. A region of interest (ROI) of approximately the size of a single nucleus was defined to standardize the measurements of nuclei between seedlings and samples. The 10 brightest nuclei from each seedling were chosen to obtain integrated density, and 8 seedlings from each independent pHYL1::HYL1-CFP line were analyzed.

### Gene Expression Analyses

Seedlings were grown in the dark as described previously for the indicated times in each experiment. ∼100 mg of collected tissue was used to extract total RNA by using the protocol described elsewhere [40]. Total RNA was then treated with DNaseI RQ1 (Promega) according to the manufacturer’s instructions. First-strand cDNA synthesis was performed using the MMLV reverse transcriptase (Invitrogen), 2 μM oligodT (dT25) and 2 μM random hexamers (Invitrogen) or 50nM stem-loop specific primers (for miRNA expression analysis). cDNA obtained was then diluted to a final concentration of ∼1 ng/μl with sterile distilled water. Real-time PCR was performed using 5 μl of cDNA, 300 nM each primer, Taq DNA Polymerase (Genbiotech, Argentina) and EvaGreen™ as dye in AriaMx Real Time PCR System (Agilent). MiRNA levels were determined by stem-loop qPCR[41], using specific forward primers for each miRNA and the reverse universal stem-loop primer (Universal SL R, S3 Table). Each PCR was repeated at least two times, and the mean expression values from three technical replicates were used for further calculations. PP2AA3 was used as a normalization control as described previously [42]. Normalized gene expression is represented relative to the 2 days’ dark-grown *hyl1-2* (in Fig 1F), *se-1* (in Fig 2E), and Col-0 (in Figs 1G, S5, 2F, 5A, 5B, 5C and 5D) set at unity. Primer sequences for qRT-PCR can be found in S3 Table.

### Protein Analyses

Total proteins were extracted using urea buffer: 50 mM Tris-HCl (pH 7.5), 4 M urea, 150 mM NaCl, 0.1 % NP-40, Protease inhibitor mix (Merck), and 50 μM MG-132 (Cayman Chemical). Protein extracts were separated by 12% SDS-PAGE, transferred to nitrocellulose membrane (Amersham), immunoblotted with a 1:1000 dilution of α-HY5 primary antibody (Agrisera) and a 1:10000 dilution of α-rabbit IgG HRP-conjugated secondary antibody (Agrisera). Chemiluminescent detection was performed with Hyperfilm ECL (Amersham). Ponceau staining was used as loading control.

### Statistical analyses

Data were analyzed by Student’s test when two sample datasets were compared or one-way ANOVA in case of more than two datasets, and the differences between means were evaluated using Tukey’s post-test (Infostat Software v. 2017, Facultad de Ciencias Agropecuarias, Universidad Nacional de Córdoba, Córdoba, Argentina). Statistically significant differences were defined as those with a P value < 0.05 or < 0.01, as indicated.

### Accession numbers

HYL1 (At1g09700), SE (At2g27100), DCL1 (At1g01040), CPL1 (At4g21670), HY5 (At5g11260), PIF3 (At1g09530), RBCS1A (At1g67090), CHS (At5g13930), PIL1 (At2g46970), XTR7 (At4g14130), CAB1 (At1g29930), MIR156b (At4g30972), MIR156c (At4g31877), MIR156d (At5g10945), MIR159b (At1g18075), MIR162a (At5g08185), MIR164c (At5g27807), MIR166a (At2g46685), MIR166b (At3g61897), MIR171b (At1g11735), PP2AA3 (At1g13320).

## Acknowledgments

We thank Diego Aguirre for plant care in growth chambers; Rodrigo Vena for technical assistance with confocal microscopy; Pablo Manavella for *cpl1-3*, *cpl1-7*, *dcl1-100*, and HYL1 phosphomimics seeds; Jaume Martínez-García for *hy5-2* seeds; Javier Botto for pHY5::HY5-YFP *hy5* seeds; Yuda Fang and David L. Spector for plasmids containing pHYL1::HYL1-CFP and pSE::YFP-SE..

## Supporting information

**S1 Fig. *Hyl1-2* mutants do not show a delay in germination.** 1-day dark-grown control (Col-0) and *hyl1-2* seedlings in 0.5 MS agarose plates.

**S2 Fig. Hypocotyl elongation in light-grown *hyl1-2* mutants.** (A) Representative *hyl1-2* and control (Col-0) seedlings grown for 3 days in long day (LD, 16 h light / 8 h dark) are shown. (B) Hypocotyl length of *hyl1-2* and Col-0 light-grown seedlings at the indicated time points. dLD indicates days in LD. Data are presented as mean ± SEM of at least 40 seedlings from 2 biological replicas. Bar in (A) = 2 mm.

**S3 Fig. Characterization of transgenic lines that express HYL1-CFP.** (A) Integrated fluorescence intensity of root nuclei from 2-days dark-grown pHYL1::HYL1-CFP *hyl1-2* seedlings. Data are presented as mean ± SEM of at least 60 nuclei from two biological replicas. AU: Arbitrary units. Statistically significant differences between groups are indicated by different letters (ANOVA, p < 0.05). (B) Representative confocal fluorescence images of roots from 2-days dark-grown pHYL1::HYL1-CFP *hyl1-2* seedlings. (C) Hypocotyl length and (D) hook angle measurements of *hyl1-2*, Col-0 and pHYL1::HYL1-CFP *hyl1-2* dark-grown seedlings at the indicated time points. Data are presented as mean ± SEM of at least 40 seedlings from 2 biological replicas. (E) pHYL1::HYL1-CFP *hyl1-2* and control plants grown for 15 days in lond day (LD).

**S4 Fig. HYL1-CFP displays two different localization patterns in darkness.** Confocal fluorescence images of roots from 3-days dark-grown pHYL1::HYL1-CFP *hyl1-2* seedlings. (A) Discrete nuclear and (B) diffuse localization patterns of fluorescence.

**S5 Fig. Expression levels of pri-miRNAs in dark-grown *hyl1-2* and *se-1* mutants.** qRT-PCR analysis of *hyl1-2*, *se-1* and Col-0 dark-grown seedlings at the indicated time points. dD, days in darkness. Expression levels of miRNA precursors were normalized to *PP2AA3* housekeeping gene and expressed relative to the Col-0 2dD value set at unity. Means ± SEM are shown from technical triplicates and 2 biological replicas. Asterisks indicate statistically different mean values compared with their corresponding wild-type (p < 0.05).

**S6 Fig. Hypocotyl elongation in light-grown *se-1* mutants.** (A) Representative *se-1* and control (Col-0) seedlings grown for 3 days in long day (LD, 16 h light / 8 h dark). (B) Hypocotyl length of *se-1* and Col-0 light-grown seedlings at the indicated time points. dLD indicates days in LD. Data are presented as mean ± SEM of at least 40 seedlings from 2 biological replicas. Bar in (A) = 2 mm.

**S7 Fig. DCL1 participates in the control of skotomorphogenesis.** Representative homozygous (*dcl1-100*) and heterozygous (*dcl1-100*/+) or control seedlings from a *dcl1-100*/+ segregating population, and control siblings (WT) grown in (A) darkness for 3 days (3dD), (B) plus 4 days in long day (3dD+4dLD). (C) Hypocotyl length and (D) hook angle measurements of *dcl1-100* and control dark-grown seedlings at the indicated time points. Data are presented as mean ± SEM of at least 10 seedlings from 2 biological replicas. Asterisks in (C) and (D) indicate statistically different mean values compared with their corresponding wild-type (p < 0.05). Bars in (A) and (B) = 5 mm.

**S8 Fig. Complementation of *hyl1-2* defects in leaves by dsRBD1 mutant versions of HYL1.** p35S::HYL1 *hyl1-2* comprising mR1 (K17A/R19A), mR2 (Δ40-46), mR3 (R67A/K68A) and wild-type (WT) mutant versions of dsRBD1, and control plants grown for 15 days in lond day (LD).

**S9 Fig. Phenotypes in leaves of *CPL1* mutant adult plants.** Mild (*cpl1-3*) and strong (*cpl1-7*) mutant alleles of *CPL1*, *hyl1-2* and control plants grown for 18 days in lond day (LD).

**S10 Fig. Epistasis analysis in *hyl1-2* and *cpl1* double mutants.** (A) Hypocotyl length and (B) hook angle measurements of Col-0, *hyl1-2*, *cpl1-3*, and double mutants *hyl1-2 cpl1-3* dark-grown seedlings at the indicated time points. (C) and (D) Representative hooks of 2- and 4-days, respectively, dark-grown mutant lines from (B). (E) Hypocotyl length and (F) hook angle measurements of Col-0, *hyl1-2*, *cpl1-7*, and double mutants *hyl1-2 cpl1-7* dark-grown seedlings at the indicated time points. (G) and (H) Representative hooks of 2- and 4-days, respectively, dark-grown mutant lines from (F). dD, days in darkness. Data are presented as means ± SEM of at least 40 seedlings from 2 biological replicas. Statistically significant differences between groups are indicated by different letters (ANOVA, p < 0.05).

**S11 Fig. Complementation of *hyl1-2* defects in leaves by HYL1 phosphorylation mimics.** pHYL1::HYL1 *hyl1-2* comprising hypophosphorylation mimics (S42A and S159A), hyperphosphorylation mimics (S42D and S159D), or wild-type (WT) version of HYL1, and control plants grown for 18 days in long day (LD).

**S12 Fig. Expression analyses of *HY5* in HY5-YFP lines.** *HY5* expression levels in 4-days dark-grown pHY5::HY5-YFP *hy5-2* (HY5-YFP) and pHY5::HY5-YFP *hyl1-2 hy5-2* (HY5-YFP hyl1-2) were measured by qRT-PCR. Expression levels were normalized to *PP2AA3* housekeeping gene and expressed relative to the HY5-YFP value. Means ± SEM are shown from technical duplicates.

**S1 Table. Localization patterns of HYL1-CFP during skotomorphogenesis.**

**S2 Table. Segregation analysis of *dcl1-100* heterozygous seedlings.**

**S3 Table. Primers used in this work.**

